# Effective Computational Framework for Pre-Interventional Planning of Peripheral Arteriovenous Malformations with In Vivo and In Vitro Validation

**DOI:** 10.1101/2021.05.15.444310

**Authors:** Gaia Franzetti, Mirko Bonfanti, Cyrus Tanade, Chung Sim Lim, Janice Tsui, George Hamilton, Vanessa Díaz-Zuccarini, Stavroula Balabani

**Author notes:** Address correspondence to Vanessa Díaz-Zuccarini and Stavroula Balabani, Department of Mechanical Engineering, University College London, Torrington Place, London WC1E 7JE, UK,. Electronic mails. Gaia Franzetti and Mirko Bonfanti have contributed equally to this work.

## Abstract

**Purpose:** Peripheral arteriovenous malformations (pAVMs) are congenital lesions characterised by abnormal high-flow, low-resistance vascular connections – constituting the so-called *nidus* – between arteries and veins. The mainstay treatment typically involves the embolisation of the nidus with embolic and sclerosant agents, however the complexity of AVMs often leads to uncertain outcomes. This study aims at developing a simple, yet effective computational framework to aid the clinical decision making around the treatment of pAVMs.

**Methods:** A computational model was developed to simulate the pre-, intra-, and post-intervention haemodynamics of an AVM. A porous medium of varying permeability was used to simulate the effect that the sclerosant has on the blood flow through the nidus. The computational model was informed by computed tomography (CT) scans and digital subtraction angiography (DSA) images, and the results were compared against clinical data and experimental results.

**Results:** The computational model was able to simulate the blood flow through the AVM throughout the intervention and predict (direct and indirect) haemodynamic changes due to the embolisation. The simulated transport of the dye in the AVM was compared against DSA time-series obtained at different intervention stages, providing confidence in the results. Moreover, experimental data obtained via a mock circulatory system involving a patient specific 3D printed phantom of the same AVM provided further validation of the simulation results.

**Conclusion:** We developed a simple computational framework to simulate AVM haemodynamics and predict the effects of the embolisation procedure. The developed model lays the foundation of a new, computationally driven treatment planning tool for AVM embolisation procedures.

## 1. Introduction

Arteriovenous Malformations (AVMs) are rare vascular developmental abnormalities, characterised by a high flow rate and low resistance shunt between arteries and veins without an intervening network of capillaries [1]. AVMs consist of feeding arteries, an intervening network of small pathological blood vessels – called *nidus* – and draining veins. The direct communication between a high-pressure arterial system and a low-pressure venous system leads to arterialised venous pressures and impairs the normal venous drainage.

AVMs can occur in every part of the body. Cerebral AVMs are associated with neurological deficits and carry a high risk of intracranial bleeding. Peripheral AVMs (pAVMs), located outside the central nervous system, are uncommon but can be extremely complex and notoriously difficult to treat. pAVMs can increase cardiac load leading to left ventricular failure, arterial insufficiency (steal) and chronic venous hypertension. Moreover, bleeding, nonhealing and painful ulcerations cause profound psychological and physical disturbances [1]. Diagnosis of AVM is based on clinical history and physical examination, supported by Doppler ultrasound, magnetic resonance imaging (MRI), computed tomography (CT) scans and intra-arterial digital subtraction angiography (DSA) [2]. DSA is a fluoroscopic technique used for visualising blood vessels in which radiopaque structures, such as bones, are “subtracted” digitally from the image, thus allowing for an accurate visualisation of the blood vessels. To image an AVM, a contrast agent (CA) is usually injected in the blood stream proximally to the AVM and its flow is imaged via X-rays.

Endovascular treatment is often regarded as the mainstay interventional therapy because it is minimally invasive, and typically involves the vascular occlusion of nidus by embolo-sclerotherapy. Embolic agents include sclerosants such as *sodium tetradecyl sulfate* (STS) foam and *absolute ethanol* (AE), metallic coils, vascular plugs, and *N-butyl cyanoacrylate*. Sclerosants induce the disruption of the endothelial cells which causes localised vasospasm and thrombosis, leading to an increase of the resistance to the blood flow through the nidus and lastly to the occlusion of the vascular lumen. Precise delivery of the embolic agent solely to the AVM nidus vasculature is key to the long-term success of the intervention and minimisation of complications. Injection of the embolic agent too proximally or distally to the AVM nidus is now thought to be futile and counterproductive as it may stimulate aggressive growth, since the nidus is able to recruit other feeding arteries and drainage veins, respectively. Embolising the proximal artery to the nidus may also prevent future access to the nidus [3]. Moreover, embolo-sclerotherapy carries the risk of disruption of the blood supply to end-organs leading to tissue ischaemia, whilst high doses of injected sclerosant may be toxic [4]. Currently, endovascular treatments are guided by multiple inter-operative DSAs. However, the limited resolution of the images, and the complex angio-architectures of pAVMs make the nidus localisation highly arduous. There are no quantitative tools to help clinicians to optimise any given treatment strategy, which relies on subjective clinical pAVM classification systems and clinical experience [5].

Numerical and experimental models of cardiovascular disease have been employed for haemodynamic studies, presurgical simulations as well as validation of imaging techniques, and they can be used as tools to support the clinical decision making around complex vascular pathologies [6]. However, due to the modelling challenges posed by AVMs, the application of these techniques to AVM management is still at its infancy.

The majority of numerical works described in the literature employed simplified approaches to model the intricate geometry of AVMs, such as the exclusive use of electric networks to reproduce the vasculature [7–11], or their combination with a porous medium to represent the nidus [11]. A similar approach was employed by Orlowski *et al*., in which the haemodynamics of cerebral AVMs were studied by means of patient-derived computational fluid dynamics (CFD). Due to difficulties in extracting the internal geometry of the nidus from the clinical images, the latter was modelled as a porous medium with parameters (i.e. permeability, porosity) calibrated based on the patient’s CT scans [12].

Experimental works aiming at reproducing *in vitro* AVM geometries have made use of similar, simplified porous media approaches to mimic the nidus, such as meshes [13], small beads in a syringe [14], or open-pore cellulose sponges [15]. Only recently, Kaneko *et al*. manufactured a phantom of a brain AVM for interventional simulations by 3D printing, and were able to reproduce the vessels of the nidus up to the limit of spatial resolution of the 3D angiograms (< 150 μm) [16].

Published computational and experimental works have successfully reproduced the patient-specific haemodynamics of AVMs, and simulated the embolisation procedure for treatment planning. However, a framework that can be generalised and implemented in near real-time to support AVM treatment planning has yet to be developed.

The present study reports on a simple, yet effective computational framework to aid the clinical decision making around the treatment of pAVM. A successful AVM embolisation procedure was simulated using CFD informed by clinical data and a simple haemodynamics framework was developed based on two hypotheses: (i) the action of the sclerosant on the nidus’ resistance to the blood flow can be modelled via a change in the permeability of the porous medium used to simulate the nidus, and (ii) a mathematical relationship can be established between the sclerosant dosage and the consequent nidus’ permeability change. The model was validated against *in vivo* DSA images and *in vitro* experimental results, demonstrating that such a computational framework can be used to predict, on a patient-specific basis, the embolic agent dosage necessary to achieve a successful embolisation, and in turn to inform and optimise the embolisation intervention.

In this work, a successful embolisation on an AVM case was considered to investigate the first of the two aforementioned hypotheses, demonstrating the feasibility of a simple haemodynamic model informed by clinical data to simulate the intervention. The model results were compared to *in vivo* DSA images and *in vitro* experimental results to demonstrate the validity of the simulation outcomes.

## 2. Materials and Methods

### Patient data and intervention

Retrospective and anonymised data of a 47-year old male patient was obtained from the Royal Free Hospital (London, UK) as part of an ethically approved protocol (NHS Health Research Authority, ref: 19/SC/0090). Anatomical data available from CT scans showed an extracranial Yakes Type III AVM fed by an arterial branch originating from the occipital artery (Figure 1a,b). The AVM was treated with two different sclerosant agents: a mixture of STS (Fibrovein 3%, STD Pharmaceutical Products, UK) and air, foamed using the Tessari method [17] in a volume ratio of 1:4, respectively, and AE. The two sclerosants were injected via fluoroscopic guided direct puncture of the nidus in four stages as shown schematically in Figure 1c. Both sclerosants were injected at a rate of approximately 1-2 ml/s, around 2-3 ml per injection.

**Fig. 1.**
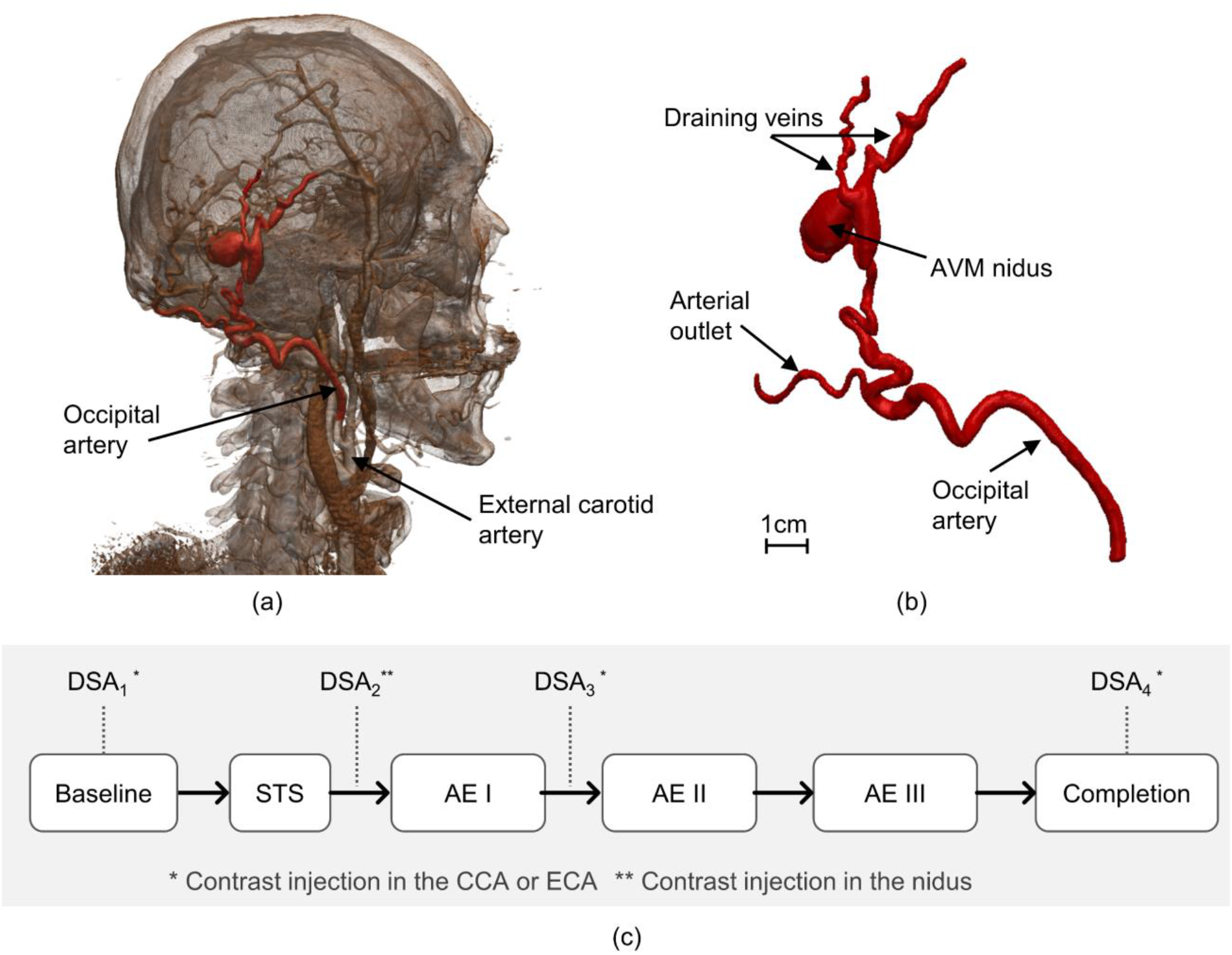
(a) 3D rendering obtained from patient’s CT scan; (b) geometry of the AVM as obtained from the segmentation procedure; (c) flowchart indicating the intervention stages and DSA images available

Several DSAs were performed to guide the intervention by injecting approximately 5-10 ml of a half-strength contrast agent (CA) (50% of Omnipaque 240 (GE Healthcare, USA) and 50% of a saline solution) in about 1-2 seconds via a catheter positioned in the common carotid artery (CCA) and external carotid artery (ECA) via the right common femoral artery (CFA) access, or via a percutaneous needle directly in the nidus (Figure 1c). DSA images were acquired with a rate of 2 frames per second.

### Image segmentation and flow rate estimation

The patient-specific AVM geometry was segmented using Simpleware ScanIP (Synopsys, USA) via semi-automated segmentation tools based on thresholding. The geometry, shown in Figure 1b, included the feeding artery of the AVM (diameter AI: ∼3.8 mm), one arterial outflow branch (diameter AO: ∼2 mm), the nidus (size: 16 × 12 × 9 mm) and two draining veins (diameter VO_1_ : ∼1.6 mm, diameter VO_2_ : ∼ 2.3 mm).

DSA images allowed the estimation of the blood flow rate in the feeding artery (Q_in_) from the ratio of the volume swept by the CA between two consecutive DSA frames, to the elapsed time between them (i.e. 0.5 s). The flowrates at three stages of the intervention (Figure 1c), baseline, Post AE I, and Post AE III, were thus obtained and found equal to 1.645 ± 0.040, 1.011, 0.231 ± 0.021 ml/s, respectively (mean value ± standard deviation (SD)). Measurements were repeated three times at different locations on the feeding artery and/or on different DSA time-series. However, enough data points were not available for the Post AE I stage, therefore SD is not provided.

### Computational model

A computational model of the blood flow and CA transport in the AVM was implemented with the commercial CFD software CFX (ANSYS, US). The blood was modelled as an incompressible Newtonian fluid with density equal to 1060 kg/m^3^ and a dynamic viscosity of 4 × 10^−3^ Pa s.

A schematic of the computational model and its boundary conditions (BCs) is shown in Figure 2. To model the blood flow, a pressure condition (P_in_) was set at the inlet to simulate the action of the heart, while zero pressure conditions were prescribed at the two venous outlets. To model the effect of the vascular bed downstream to the arterial branch, a resistance (R_art_) was coupled to the arterial outlet by setting the following relationship between the outlet mean pressure (P_art_) and the flow rate (Q_art_):

**Fig. 2.**
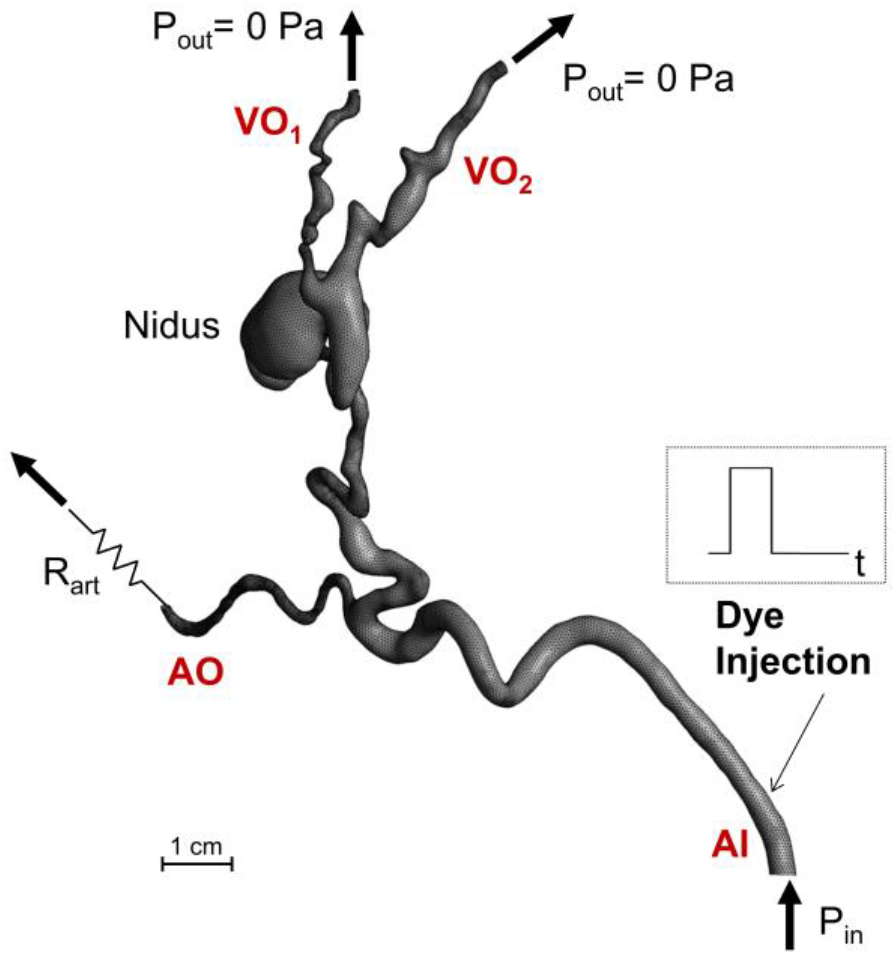
Schematic of the numerical model and its boundary conditions. AI, arterial inlet; AO, arterial outlet; VO, venous outlet

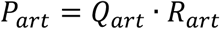

P_in_ and R_art_ were calibrated to match the baseline DSA-derived flow rate value and the CA distribution among the outlet branches. The obtained P_in_ and R_art_ values, equal to 20.837 mmHg and 24.138 mmHg s/ml, respectively, were kept constant for all the simulations.

A porous medium approach was adopted to model the resistance the blood experiences when flowing through the nidus as a result of the injection of the sclerosant. To this end, an isotropic momentum loss – expressing the Darcy’s equation [18] - was prescribed to the nidus region by adding the momentum source **S**_M_ [N/m^3^] to the right-hand-side of the Navier Stokes equations:

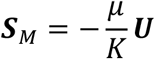

where μ is the blood dynamic viscosity, **U** is the velocity vector and K [m^2^] is the permeability. K was tuned for each intervention stage to match the CA transport in the AVM as observed in the DSA images. For the baseline model, K was set equal to 2 × 10^−8^ m^2^ which translates into a near-zero resistance to the blood flow (i.e. nidus pressure drop = 0.4 mmHg), while for the completion model, K was decreased until almost no flow was obtained in the nidus (i.e. Q_nidus_ ≤ 1% of Q_in_), as observed in the DSA images. Clinical DSA images were not available for the Post AE II intervention stage, therefore the value of K for the corresponding model was obtained via a quadratic interpolation between the known K values calibrated for the Post STS, Post AE II and Post AE III stages.

To simulate the CA transport through the AVM, the following convection-diffusion equation was solved in the model domain:

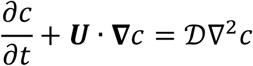

where *c* [kg/m^3^] is concentration per unit mass of the contrast agent and 𝒟[m^2^/s] is its kinematic diffusivity, equal to 2.2 × 10^−10^ m^2^/s [19]. An initial condition of *c* equal to zero was set in the domain. A zero-flux condition was prescribed at the walls, while the concentration *c* at the inlet was set equal to a unity-amplitude square wave of 1.5 s width to simulate the CA injection (Figure 2). Thus, the inlet concentration *c* was set equal to 1 kg/m^3^ for 1.5 s, and to zero for the remaining time of the simulation, in agreement with the available clinical information. The simulated total time was equal to 5 s to allow CA to travel throughout the domain and the comparison of the simulation results against the DSA images.

### Mesh and sensitivity

The geometry was meshed with Fluent Meshing (ANSYS, USA) using a tetrahedral mesh in the core region (minimum and maximum element size equal to 0.1 and 0.8 mm, respectively) and 7 inflation layers at the walls to accurately resolve the boundary layer (first layer height = 0.02 mm, growth rate = 1.2) resulting in a computational grid of approximately 860K elements. A finer mesh of 3M elements was also generated for the mesh sensitivity analysis. Mesh independence was demonstrated by comparing the outlet flow rates of the two meshes, resulting in a discrepancy of less than 1.5%. The medium mesh was therefore adopted for this study.

### Numerical simulations and post-processing

The fluid dynamics and CA transport equations were solved in a de-coupled fashion to decrease the computational time with the commercial finite-volume solver ANSYS CFX v18. First, the steady-state Navier Stokes and continuity equations were solved using a high-resolution advection scheme. The flow was assumed laminar (Reynolds number at the inlet equal to 150) and the wall of the AVM domain rigid. Due to the low acquisition rate of the DSA data, only an estimate of the average blood flow in the AVM was available to inform the haemodynamic models, which were therefore assumed as steady. The velocity fields obtained from the haemodynamic models were subsequently input to the time-dependent transport equation of the dye, which was solved using a high-resolution advection scheme and a second order backward Euler scheme with a fixed time step of 0.01 s. A target maximum RMS residual of 10^−5^ was set to guarantee the convergence of the solution.

### Experimental setup

A rigid replica of the AVM was manufactured in a clear resin via stereolithography 3D printing (Formlabs, USA) in a scale of 2:1 (Figure 3a). The phantom was connected to an experimental setup (Figure 3b) aimed at reproducing the computational BCs and demonstrate the feasibility of *in vitro* testing for AVM treatment planning. Water was used as a working fluid. A plastic mesh was employed as porous medium in the nidus to simulate the sclerosant-induced increased resistance. The permeability value was decreased by increasing the mesh density.

**Fig. 3.**
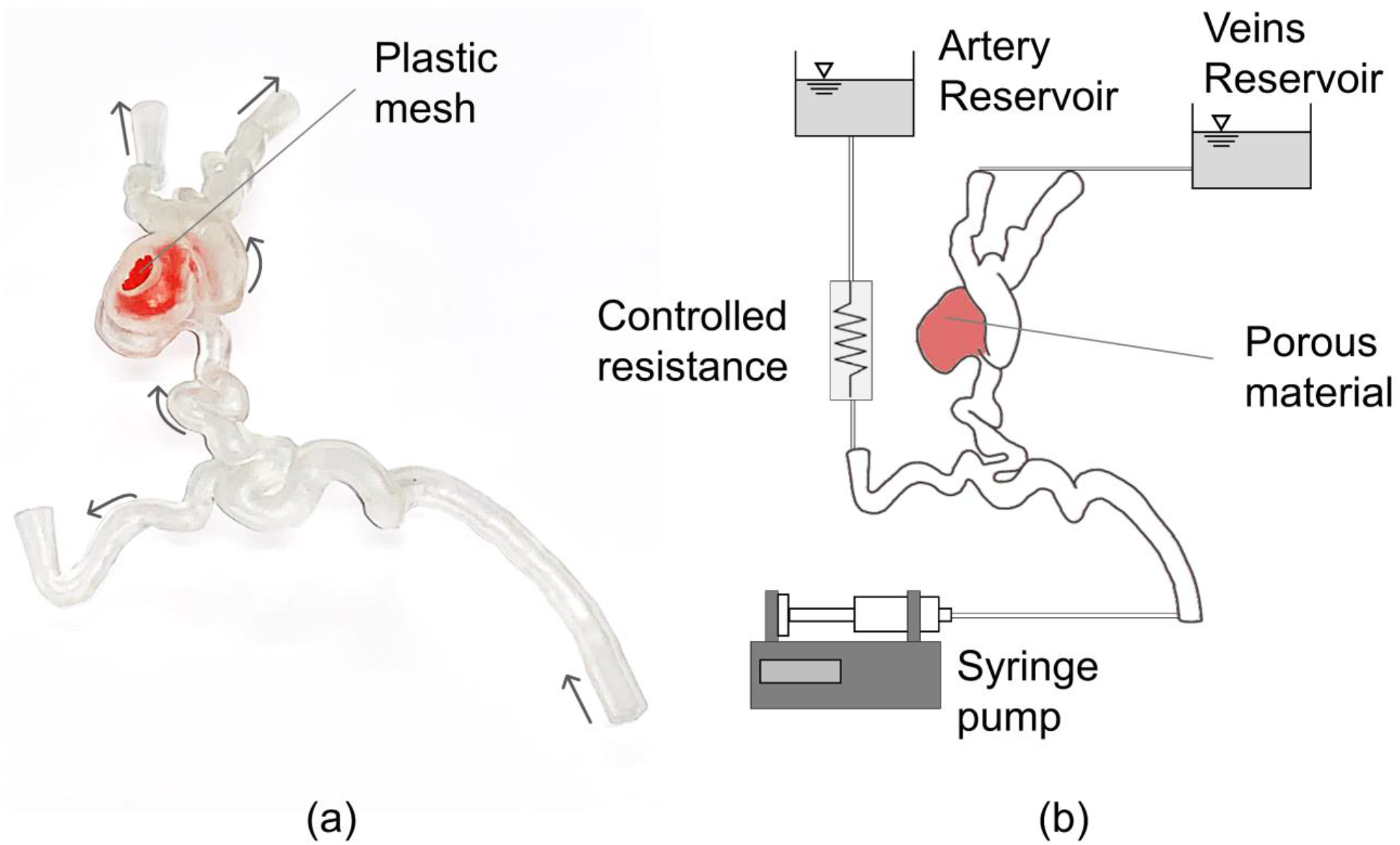
(a) Picture of the 3D printed AVM phantom, including the plastic mesh (in colour red) employed to simulate the porous material in the nidus to reproduce the injection-induced decrease in permeability. Arrows indicate the flow direction. (b) Schematic of the experimental set up

A controlled-volume of dye (water-based food colouring, PME, UK, mixed with water) – simulating the CA – was injected upstream of the AVM model with a syringe pump (Chemyx Fusion 4000). To obtain similarity for the transport of the dye between the computational model - which mimics the *in vivo* scenario - and the experiment, the flow rates prescribed at the inlet of the phantom were double the corresponding computational values. Therefore, the numerical and experimental Péclet numbers (Pe) in the baseline stage (where *Pe* = *Dv*/𝒟, with D the inlet diameter, *v* the inlet velocity and 𝒟the diffusion coefficient, equal to 2 × 10^−10^ m^2^/s for the experimental model [20]) were equal to 2.6 × 10^6^ and 2.8 × 10^6^, respectively. Video recordings (4K resolution at 60 fps) of the experiments were acquired to capture the distribution of the dye for each phase of the treatment.

## 3. Results and discussion

### AVM haemodynamics

The computational model allowed the simulation of the blood flow within the AVM and the haemodynamic changes induced by the sclerotherapy in a simplified, but effective way.

First, it was possible to simulate the baseline stage in line with the available clinical data. Figure 4 shows the pressure distribution and streamlines obtained in this setting. Pressure drops of 11 and 21 mmHg were obtained between the inlet and the arterial and venous outlets, respectively. The lower pressure drop predicted between the inlet and the arterial branch is due to the resistance coupled to this outlet, which simulates the downstream vasculature not included in the 3D geometry. The ratio between the arterial and venous outflows was equal to 40%, which is explained by the low resistance of the nidus that causes most of the blood to bypass the capillary bed downstream of the arterial outlet by flowing through the AVM.

**Fig. 4.**
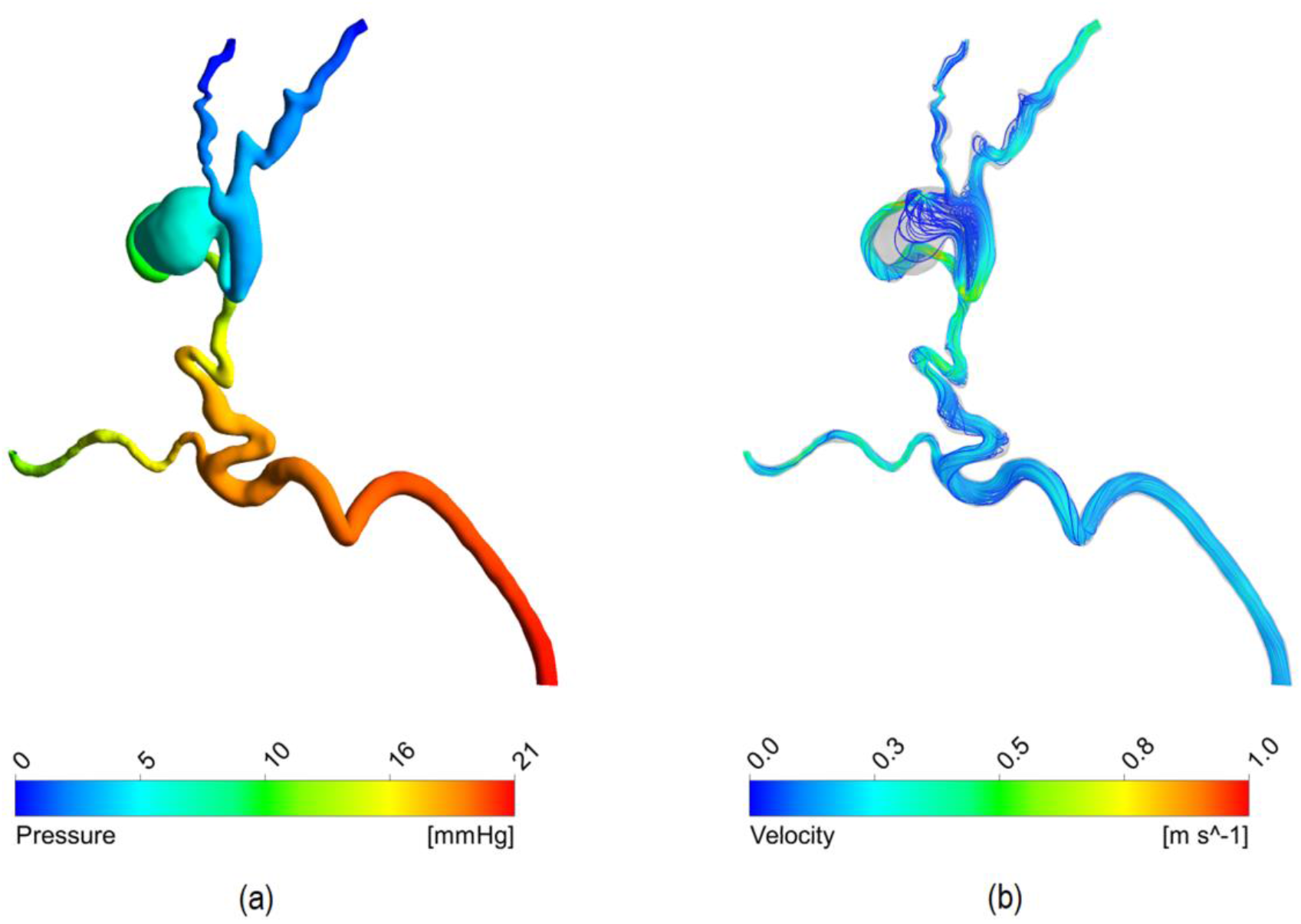
Haemodynamic results of the baseline simulation: (a) pressure distribution, (b) streamlines

According to the clinical data, the baseline inlet flow rate was equal to 1.645 ml/s. This corresponds to approximately 40-50% of the ECA flow rate – approximately 3-4 ml/s [21] – and it is higher than expected in normal physiological conditions due to the low resistance/high flow rate shunt characterising the AVM. In fact, given that the occipital artery is one of four or five branches out of the ECA, the expected flow rate ratio should be around 20-25%.

The haemodynamic changes induced by the sclerotherapy intervention were simulated by reducing the permeability *K* of the nidus (Figure 5). The primary effect following the reduction of *K* is a decrease of the Q_nidus_. As shown in Figure 5a, Q_nidus_ gradually decreased with each sclerosant injection from 1.168 to 0.005 ml/s, which corresponds to 71 and 1% of the inlet flow rate, respectively. Secondary effects include a decrease of the inlet flow rate (Figure 5b) and an increase of the flow rate through the arterial outflow (Figure 5c).

**Fig. 5.**
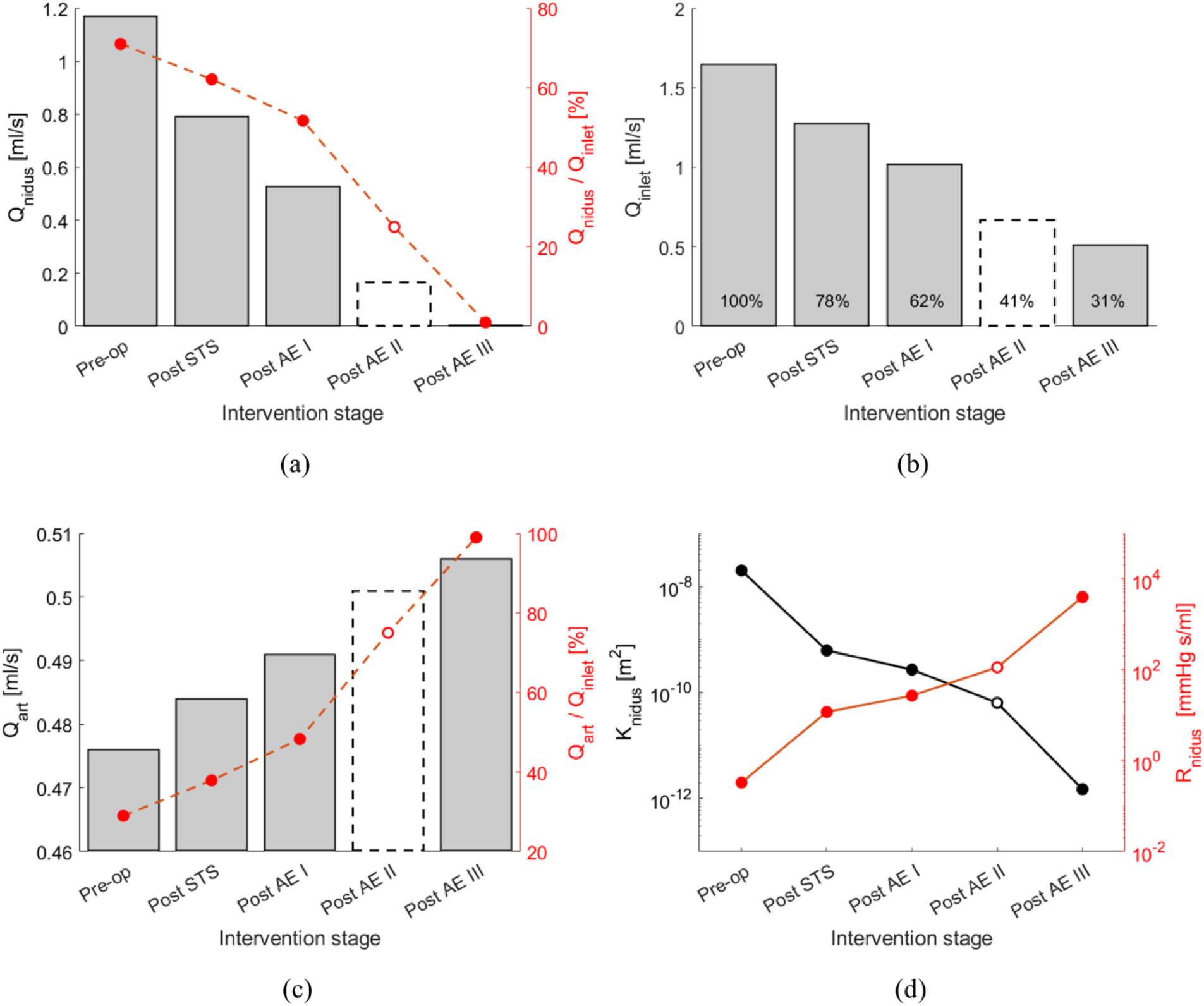
(a, b, c) The charts show the haemodynamic changes predicted by the computational model at the different stages of the sclerotherapy intervention in terms of (a) flow rate through the nidus (Q_nidus_), (b) inlet flow rate (Q_inlet_), (c) flow rate via the arterial outlet (Q_art_). (d) Nidus permeability (K) and corresponding hydraulic resistance (R_nidus_) at the different intervention stages. The results for the Post AE II stage are reported with a dashed line because, due to the lack of DSA images, the Post AE II model parameters were estimated via interpolation of the other stages rather than calibrated with clinical data

The model predicted a reduction of the inlet flow rate to 31% of the initial flow. This value is higher than what observed in the clinical data (14%) and it could be explained by post-intervention adaptations in the vascular system (e.g. a decrease of central blood pressure, an increase of the peripheral resistance) not simulated in the model.

As shown in Figure 5c, the arterial outlet flow rate increases with each sclerosant injection up to 0.5 ml/s in the completion stage, which corresponds to an increase of 6% compared to the baseline setting. The haemodynamic changes described lean towards the restoration of the physiological flow, characterised by a lower workload on the left ventricle and a higher blood flow towards the capillary beds.

Figure 5d reports how the permeability *K* of the nidus decreased during the intervention from 2 × 10^−8^ to 1.5 × 10^−12^ m^2^, which corresponds to an increase of the nidus hydraulic resistance from 0.330 to 3970 mmHg s ml^-1^.

### Contrast Agent transport in the AVM

Figure 6 shows the comparison between the transport of the CA within the AVM at 4 successive instants acquired via DSA and the corresponding simulation results at the baseline, post-EA I and post-EA III (i.e. completion) stages. A very good agreement can be noted between the clinical and computation results. In the baseline phase, the numerical results show the dye distributed amongst the nidus – and venous branches – and the arterial outlet. The comparison of the images in the post-AE I phase, shows less flow volume reaching the nidus and perfusing the venous side as expected due to the higher nidus resistance induced by the first two sclerosant injections. Lastly, Post AE III, that represents the completion stage, shows that the nidus is completely bypassed, and the inlet flow only reaches the arterial branch providing evidence of a successful intervention. The excellent agreement between the computational results and clinical data provides confidence on the suitability of representing the sclerosant action through a decreased nidus permeability, and on the accuracy of its values in the different operative stages.

**Fig. 6.**
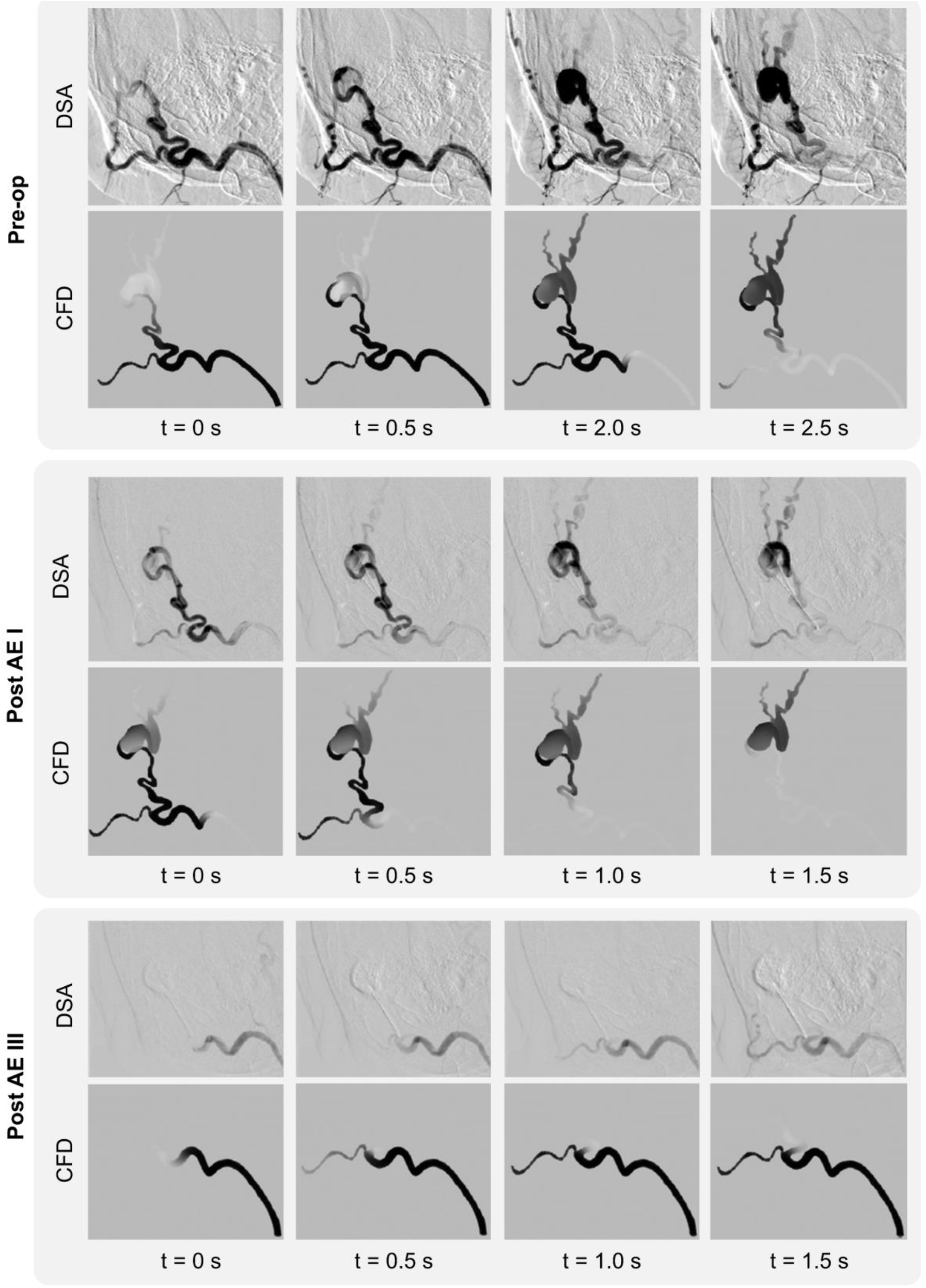
Comparison between the DSA images and CFD results showing the contrast agent transport in the AVM at four successive time instants at the Pre-op, Post AE I and Post AE II stages

### Experimental model

A simple experimental model was used to replicate *in vitro* the AVM fluid dynamics and embolisation effects simulated *in silico*.

The 3D printed phantom successfully reproduced the geometry of the case study under consideration (scale 2:1) and the clear resin used in the stereolithography process ensured sufficient transparency for flow visualisation. By altering the density of the mesh located in the nidus, it was possible to control its permeability according to the different stages of the intervention. A good comparison between computational and experimental dye distributions within the AVM was obtained as shown in Figure 7 for two selected instants in each of the three interventional stages (i.e. baseline, Post STS and Post AE III).

**Fig. 7.**
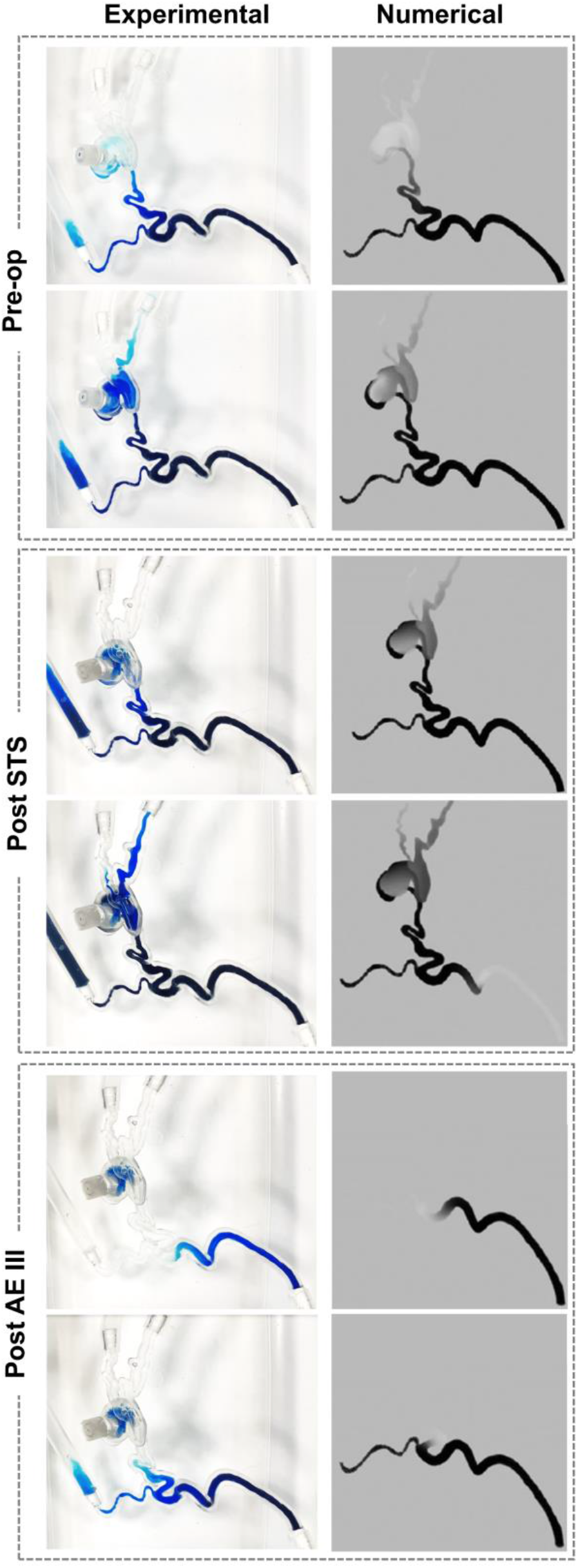
Comparison of numerical predictions and experimental results simulating the contrast agent transport in the AVM model in the baseline, Post STS and Post-AE III stages

### Clinical Application

Sclerosant agents are associated with complications and morbidity and their safe use in AVM embolo-sclerotherapy procedures is essential. The intervention requires precise delivery into the nidus, and injections too proximal to the feeding arteries should be avoided as it may lead to incomplete occlusion of the nidus, and undesired end-tissue ischaemia [1]. Currently, due to the extremely complex and patient-specific anatomy of some AVMs, these interventions usually require multiple sclerosant injection stages, guided using contrast agents and DSA (Figure 8a). Sub-optimal outcomes are frequent [2].

**Fig. 8.**
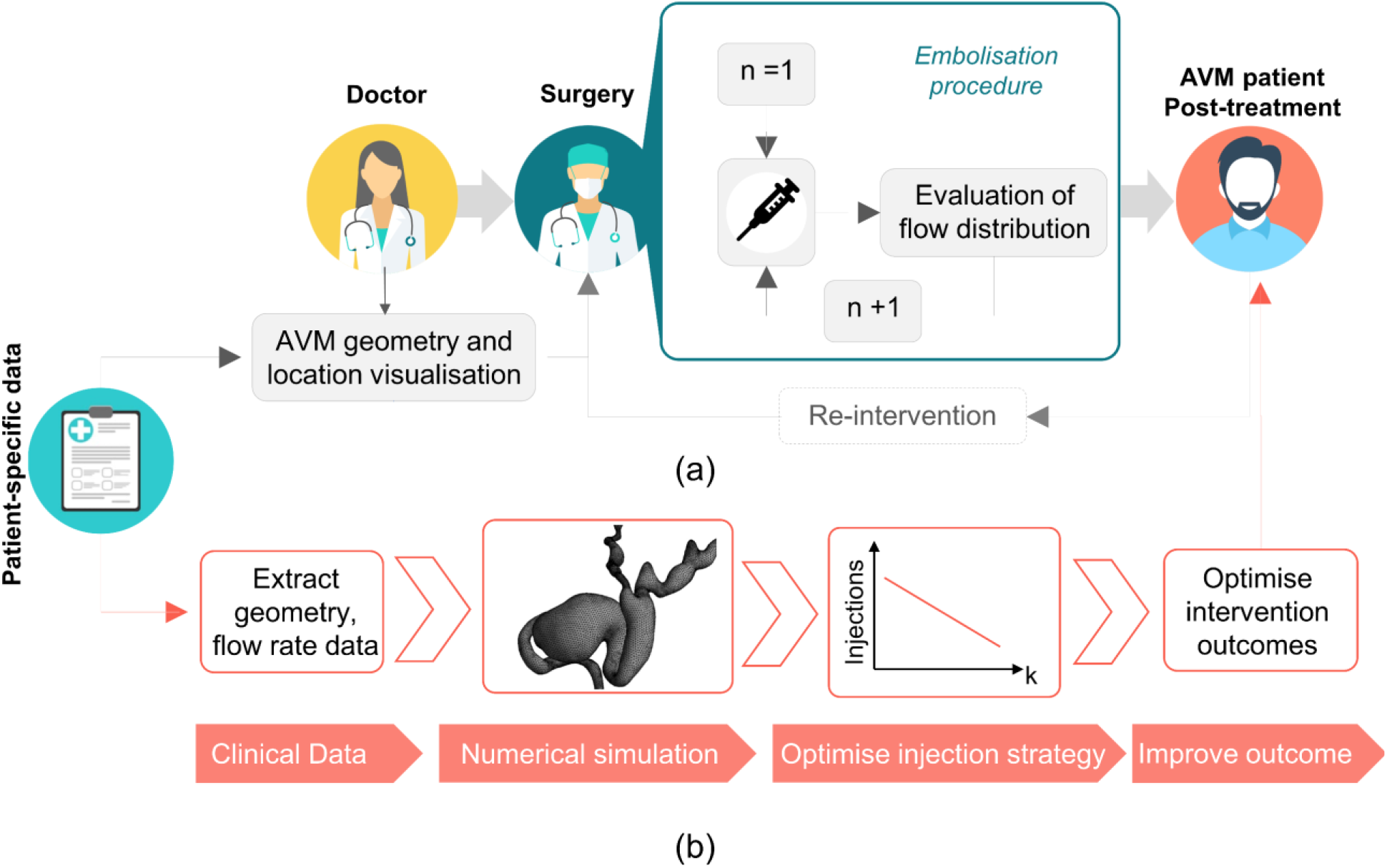
Simplified schematic of (a) the current clinical pathway for AVM interventions via embolization procedure, and (b) proposed pathway for optimised interventions planned via a computational model

In this context, a pre-interventional planning tool able to simulate the patient-specific haemodynamics of AVM and optimise the embolisation intervention would be of outmost clinical value. In this study, using computational flow dynamics and experimental validation, we were able to quantify the pre- and intra-intervention AVM haemodynamics and the effects of the sclerosant on the nidus permeability. A clear link between nidus permeability and embolisation success was demonstrated, proposing thus an optimised, computationally driven AVM treatment pathway as illustrated in Figure 8b. First, clinical data (e.g. CT or MRI scans for the geometry and doppler ultrasound or PC-MRI for the haemodynamics) are collected from the patient and, second, are used to set up a patient-specific computational model able to predict the nidus permeability *K* necessary to satisfactorily reduce the blood flow through the AVM and restore the physiological haemodynamics. The optimal nidus permeability change is then used to estimate the right sclerosant dosage needed for a successful intervention.

Future studies are needed to explore the possibility of establishing a mathematical relationship between the sclerosant dosage and the corresponding change in *K*. With this aim, the computational framework developed in this study will be applied to a cohort of patients undergoing embolisation procedures, and a mathematical relationship describing the correlation between the sclerosant dosage and simulated *K* change will be evaluated.

### Limitations

The work was developed with the goal of supporting pre-interventional planning of pAVMs by using resources available in a clinical setting. Therefore, only CT and DSA images were used to develop and inform the developed models. However, deriving flow information from DSA data is subject to uncertainty. In future work, Doppler ultrasound and/or PC MRI will be used to acquire flow rate data.

The models were developed to be as simple as possible to allow for almost-real time simulations that can be implemented easily and in a timely manner. Therefore, continuous flow, rigid walls and an ‘empirical’ modelling of the sclerosant action were employed. Despite these approximations, the excellent comparison with DSA data indicates that the simplified model developed within this work is an effective way to represent the macroscale behaviour of the interventions. Lastly, the case study considered in this works represents a simple case of pAVM, with a limited number of feeding arteries and draining veins, and a well-located nidus. More complex AVM anatomies would require further assumptions. For instance, the intricate vessels and micro vessels forming the AVM nidus could be modelled via a porous medium with spatially-varying CT-calibrated parameters as described in Orlowski *et al*. [12].

## 4. Conclusion

In this work, a patient-specific haemodynamic model was implemented to simulate the embolisation procedure of a peripheral AVM. The model was developed to verify our hypothesis that the haemodynamic effects of the embolisation procedure can be simulated via a porous medium with varying permeability. The hypothesis was successfully verified via the application of the computational framework to a case study informed by clinical data, and the obtained results were successfully validated via comparisons against DSA and experimental data.

The computational model developed is a fundamental building block of the proposed treatment-planning framework for AVM embolisation. Future work will be carried out to test the computational model on a cohort of AVM patients, and to establish a correlation between the nidus permeability change and the injected sclerosant dose. If successful, the proposed framework will allow the optimisation of embolisation procedures via the quantitative estimation of the correct dosage of sclerosant needed for a successful intervention, therefore decreasing the risk of complications and improving the outcome.

## Acknowledgements

The authors would like to thank Prof. Vivek Muthurangu (Great Ormond Street Hospital for Children, London, UK) for the insightful discussions, and are grateful to The Butterfly AVM Charity (butterflyavmcharity.org.uk) for their support.

## Ethics approval

The study was ethically approved protocol (NHS Health Research Authority, ref: 19/SC/0090).

